# Controllability of molecular pathways

**DOI:** 10.1101/560375

**Authors:** Stefan Wuchty

## Abstract

Inputs to molecular pathways that are the backbone of cellular activity drive the cell to certain outcomes and phenotypes. Here, we investigated proteins that topologically controlled different human pathways represented as independent molecular interaction networks, suggesting that a minority of proteins control a high number of pathways and *vice versa*. Transcending different topological levels, proteins that controlled a large number of pathways also controlled a network of interactions when all pathways were combined. Furthermore, control proteins that were robust when interactions were rewired or inverted also increasingly controlled an increasing number of pathways. As for functional characteristics, such control proteins were enriched with regulatory and signaling genes, disease genes and drug targets. Focusing on evolutionary characteristics, proteins that controlled different pathways had a penchant to be evolutionarily conserved as equal counterparts in other organisms, indicating the fundamental role that control analysis of pathways plays for our understanding of regulation, disease and evolution.

## INTRODUCTION

Modern network research recently started to focus on the development of different methods to find nodes that control entire networks (Ishitsuka et al, 2016; Liu et al, 2011; Nacher & Akutsu, 2012; Nacher & Akutsu, 2014; Vinayagam et al, 2016). Nodes that allow the topological control of underlying biological networks were found important for different cellular processes (Basler et al, 2016; Ishitsuka et al, 2016; Vinayagam et al, 2016; Wuchty, 2014; Wuchty et al, 2017). A recent analysis of a directed protein-protein interaction network indicated the presence of control proteins that were enriched with disease genes and drug targets as well as carried genomic alterations in diverse cancer types (Vinayagam et al, 2016). While these results were found in a single interaction network, the inner workings of a cell are usually organized through an elaborate network of distinct molecular pathways. In particular, each pathway is represented as a network of directed molecular interactions that provide a certain cellular function. As a consequence of the representation of pathways as directed networks, we surmised that pathway-specific proteins may allow the control of a given pathway. As pathway crosstalk is established through proteins that appear in more than one pathway, we hypothesized that sets of proteins may exist, controlling many different pathways at the same time. As a consequence, proteins that control many different pathways may mediate functional, biomedical and evolutionary significance, indicating *e.g.* disease or essential genes.

To fill this knowledge gap, we determined and analyzed proteins that structurally controlled pathways, represented as separate directed networks of interactions between proteins. Notably, we found that a small minority of proteins controlled a high number of pathways and *vice versa*. Transcending different topological levels, proteins that increasingly controlled pathways also appeared as control proteins in a combined pathway network that we obtained from pooling interactions from all underlying pathways. Furthermore, proteins that controlled a large number of pathways appeared to be resilient to rewiring and flipping the direction of interactions. Strongly indicating their biological significance, control proteins were enriched with regulatory and signaling genes, disease genes and drug targets on both topological levels. Anticipating that such topological features may carry an evolutionary blueprint we also observed that proteins that controlled different pathways had a penchant to be evolutionarily conserved as equal control counterparts in other organisms.

## RESULTS

As the majority of pathway specific interactions were directed, indicating flow of biological information from *e.g.* a transcription factor to an expressed gene, we utilized 276 human KEGG (Ogata et al, 1999) pathways that had at least 5 directed interactions. In each pathway, we mapped directed interactions to a bipartite graph, where partitions referred to proteins that started and ended direct interactions. To find potential control proteins, we determined the largest subset of interactions in each pathway network called a maximum matching, where no two interactions shared a common start and end point. Unmatched nodes that correspond to any maximum matching were previously shown that they can be chosen as a driver nodes to structurally control the whole underlying network (Liu et al, 2011). To further assess the relevance of nodes we considered the topological consequences of their removal. In particular, we defined a node as a control node, if the total number of driver nodes increased in a maximum matching after its removal (Vinayagam et al, 2016). Such nodes are considered important for the control of the underlying network as more driver nodes emerged as a consequence of their removal (**Fig 1A**). Determining such control proteins in each pathway specific network separately we counted the number of pathways that a protein controlled. Notably, we observed that the corresponding frequency distribution followed a power-law like decay (**Fig 1B**), indicating a small minority of proteins that controlled a high number of pathways and *vice versa*. To show the independence of our results from the underlying pathway data, we determined control proteins in 1,192 human Reactome (Jupe et al, 2012) pathways and corroborated our initial finding (**Fig EV1A**). Furthermore, we hypothesized that proteins that controlled an increasing number of different pathways separately may also appear as control proteins in a network that was composed of all pathway interactions. Pooling all 276 KEGG pathways, we obtained a network of 67,038 directed interactions between 5,398 proteins and found 577 (10.7%) control proteins. More quantitatively, we randomly sampled such a set of control proteins in the combined pathway network and determined their enrichment in bins of proteins that controlled an increasing number of KEGG pathways. In **Fig 1C**, we observed that these proteins were enriched in groups of proteins that controlled an increasing number of pathways while non-control proteins appeared diluted. To establish the independence of our results from the choice of pathway data, we pooled all 1,192 Reactome pathway specific networks, obtaining a directed network of 180,020 edges 8,084 proteins. Randomly sampling all 941 (11.6%) control proteins we observed similar results in the pooled network of interactions in Reactome pathways (**Fig EV1B**).

**Figure 1.**
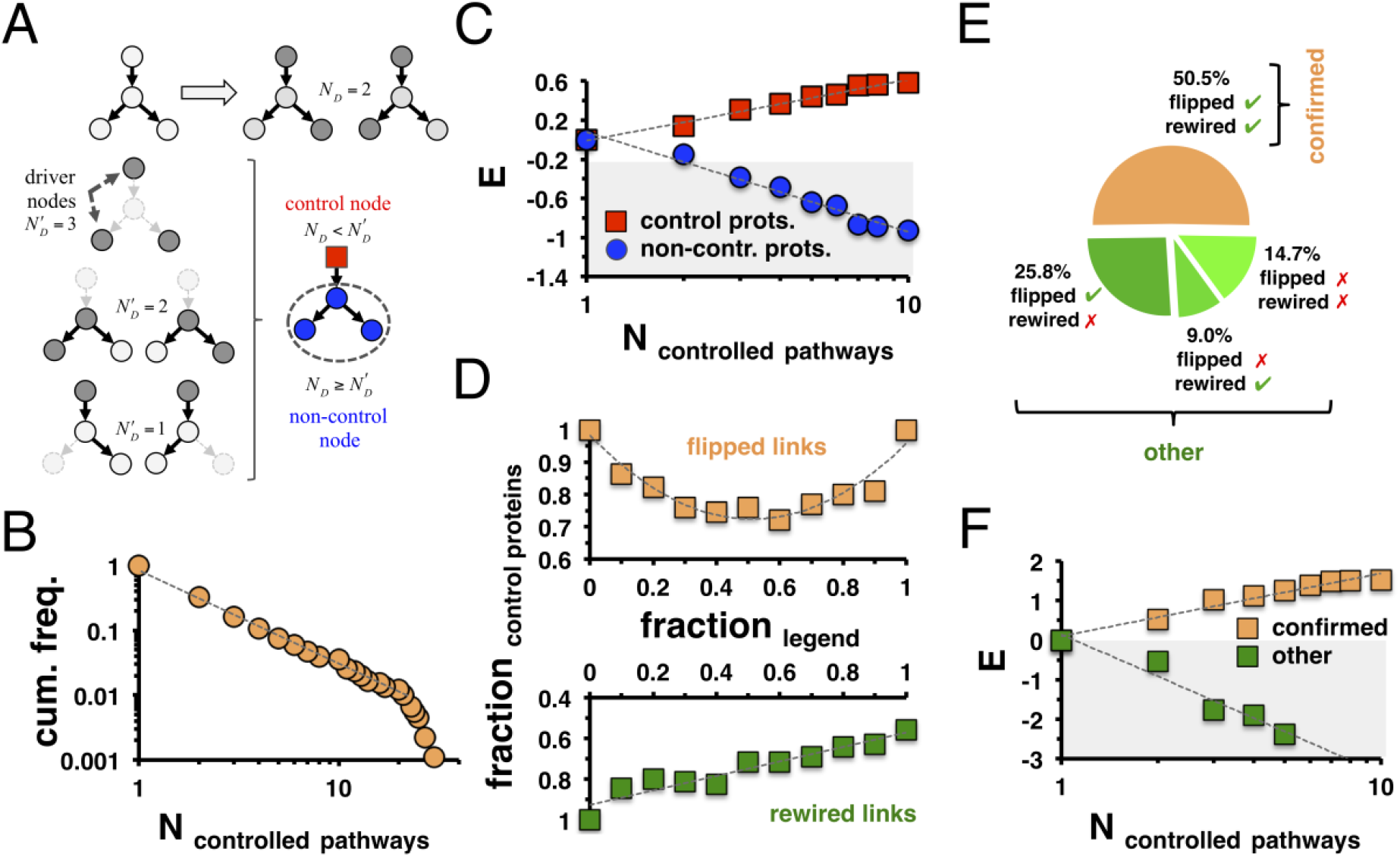
Topological characteristics of pathway controlling proteins. **A** In the schematic representation of the controllability framework the application of a maximum matching algorithm allows the determination of *N*_*D*_ =2 driver nodes. To find control nodes, we separately eliminated each node and determined the number of driver nodes 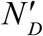 in the network thus obtained. We found a control node if the elimination of a node increased the number of driver nodes compared to the unperturbed network, 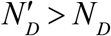. **B** Utilizing 276 KEGG pathways, we found that the cumulative frequency distribution of the number of pathways that proteins controlled followed a power-law, suggesting that a minority of proteins controls a large number of pathways and *vice versa*. **C** After determining control nodes in a network of 67,038 directed interactions between 5,398 proteins that we obtained by combining interactions of all pathways we randomly sampled such sets of control proteins. Notably, control proteins in the combined interaction network were enriched in groups of proteins that controlled an increasing number of pathways. **D** In the combined network we flipped and rewired given fractions of interactions. Notably, flipping the direction of roughly half of all interactions limited our ability to confirm control proteins the most. In turn, rewiring interactions continuously decreased the fraction of confirmed nodes. **E** When we flipped 50% of all interactions and rewired all interactions, respectively, half of all control proteins were confirmed. **F** More quantitatively, we randomly sampled sets of control proteins that were confirmed after flipping and rewiring interactions and found that such proteins were enriched in groups of proteins that controlled an increasing number of pathways. In turn, the set of remaining proteins was found strongly diluted in groups of proteins that controlled a limited number of pathways.

In a robustness analysis, we flipped the direction of given fractions of interactions in the combined pathway network and determined the number of control nodes in a network thus obtained (upper panel, **Fig.1D**). Corroborating earlier results in a network of directed protein-protein interactions (Vinayagam et al, 2016), flipping half of all interactions corresponding to the lowest fraction of control nodes that were found in the original, unperturbed network. Furthermore, we rewired a given fraction of directed interactions keeping the underlying degree distributions of nodes in the unperturbed network, indicating that half of all control nodes were robust toward the complete rewiring of the network (lower panel, **Fig 1D**). To paint a coherent picture of the robustness of control proteins, we determined control nodes in a network where we flipped one half of all directed interactions. Furthermore, we determined control nodes in a network where we rewired all interactions, keeping the degree distributions of the underlying nodes. **Fig 1E** indicates that 50.5% of all control nodes in the underlying unperturbed network were robust in the presence of rewired and flipped interactions. More quantitatively, we randomly sampled sets of control proteins that were confirmed after flipping and rewiring interactions in **Fig 1F**. Notably, such robust control proteins were almost entirely found enriched in bins of proteins that controlled a large number of pathways. To corroborate our results, we repeated this analysis using the combined network of Reactome pathways and observed similar results (**Fig EV1C-E**).

On a functional level, we presented the 20 most KEGG pathway-controlling proteins in the table of **Fig 2A** and observed that they were frequently essential for the survival of the cell. While these proteins were hardly transcription factors and membrane bound receptors, they also frequently carried kinase and signaling functions when we excluded membrane bound proteins (**Fig 2A**). On a more quantitative level, we randomly sampled 2,708 essential human genes (Chen et al, 2012; Luo et al, 2014) that we found enriched among proteins that controlled an increasing number of pathways (**Fig 2B**). Focusing on control proteins in the combined pathway network, we found that essential genes were significantly enriched as well, while they appeared diluted in the set of remaining proteins (**Fig EV2A**). Considering 4,408 proteins that were involved in signaling functions (excluding membrane bound proteins), we found that such proteins appeared enriched, while 5,701 receptor proteins appeared diluted among proteins that control an increasing number of pathways (**Fig 2C**). We obtained similar results, when we considered their enrichment in the set of control proteins in the combined network (**Fig EV2B**). As a corollary of our observation that control proteins are enriched with signaling functions, we hypothesized that such proteins may be significantly involved in regulatory processes. Considering a set of 1,471 manually curated sequence-specific DNA-binding transcription factors (Vaquerizas et al, 2009; Wilson et al, 2008) and 501 kinases (Cheng et al, 2014), we observed that proteins that control an increasing number of different pathways were more frequently enriched with kinases than transcription factors (**Fig 2D**), results that we confirmed in the combined pathway network as well (**Fig EV2C**). In turn, proteins that received post-translational modifications as a consequence of regulatory activity may be important for pathway control. Randomizing sets of methylated, acetylated and phosphorylated proteins we observed that methylated and acetylated proteins increasingly appeared in sets of proteins that controlled an elevated number of pathways while phosphorylated target appeared significantly less enriched (**Fig 2E**). Still, all targets of posttranslational modifications appeared significantly enriched in the set of proteins that controlled the combined pathway network (**Fig EV2D**). To confirm that our results were independent of KEGG pathway data, we used Reactome pathway data, indicating similar results when we considered proteins that controlled single pathways (**Fig EV3**) as well as a network of all pathway interactions combined (**Fig EV4)**.

**Figure 2.**
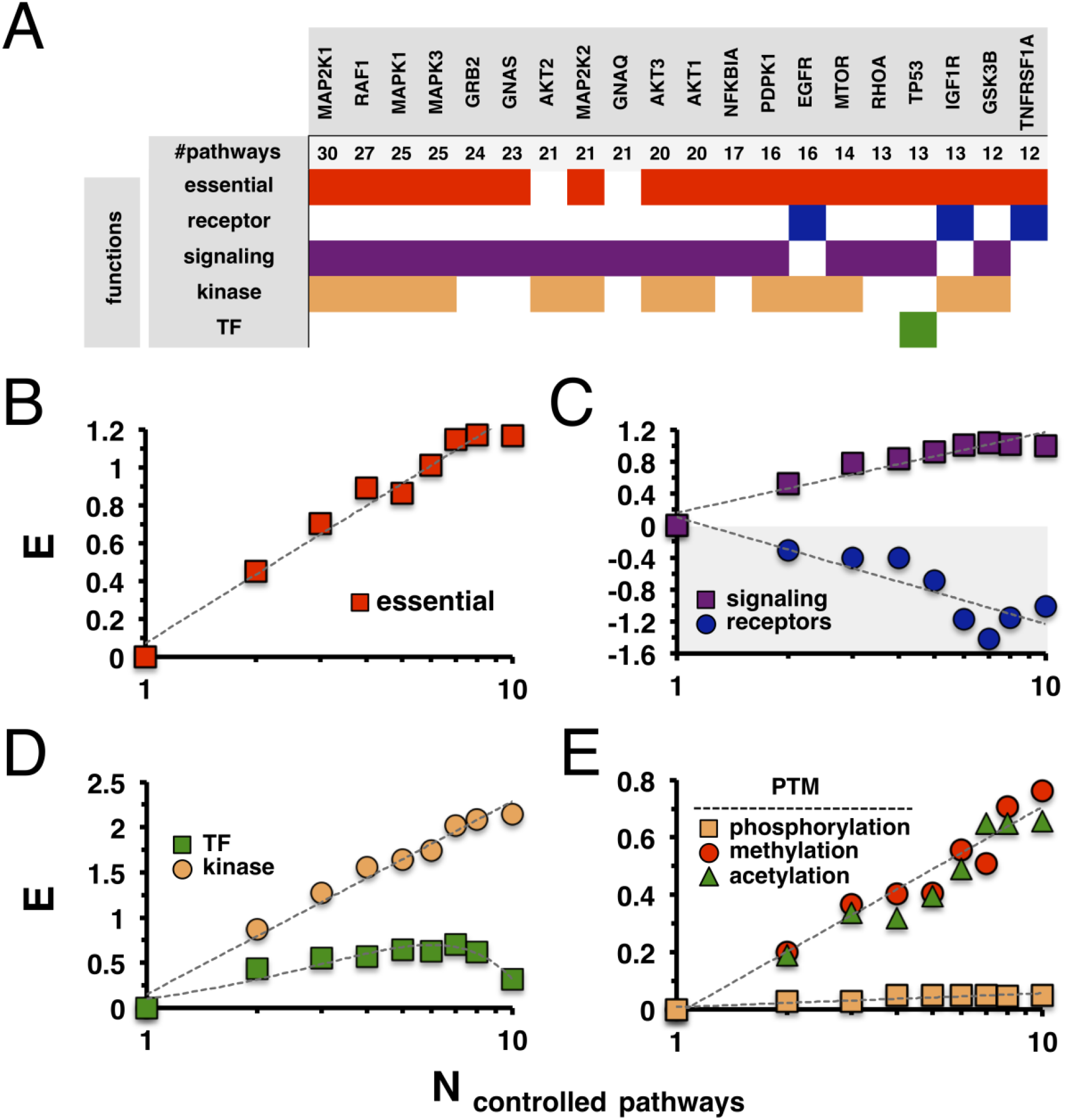
Functional characteristics of pathway controlling proteins. **A** We collected 20 proteins that controlled the highest number of pathways. We observed that such proteins were mostly essential and had kinase but rarely transcription factor functions. While sporadically membrane-bound receptors, the majority of such proteins were involved in signaling activities. **B, C** More quantitatively, we randomly sampled sets of control proteins and found that controlled an increasing number of pathways were enriched with (B) essential genes and (C) signaling functions, while they rarely were membrane-bound receptors. **D** Such proteins were more frequently enriched with kinases than transcription factors. **E** As for post-translational modifications, proteins that controlled an increasing number of pathways were strongly enriched with acetylated and methylated proteins while we only found a modest enrichment of phosphorylated proteins.

As a corollary of our previous results, we hypothesized that control proteins may play a role for the transition between healthy and disease conditions. Utilizing a set of 568 genes that were annotated by the Sanger Center as causally implicated in oncogenesis (Futreal et al, 2004), we observed that proteins that controlled an increasing number of pathways were enriched with such cancer genes (**Fig 3A**), a result that was confirmed in the combined KEGG pathway network (**Fig EV5A**). To further substantiate our observations, we considered a set of 1,259 onco- and tumorsuppressor genes that were predicted as cancer-related (Higgins et al, 2007) and obtained similar results (**Fig 3A**, **Fig EV5A**). As for viral infections, we analyzed a set of 544 human proteins that physically interacted with proteins of the HIV virus (Ako-Adjei et al, 2015) as well as 788 human genes that were down-regulated and 1,118 genes that were up-regulated upon HIV infection (Ako-Adjei et al, 2015). Notably, such infection relevant genes appeared enriched in groups of proteins that controlled an increasing number of pathways (**Fig 3B**), a result that was corroborated with proteins that controlled the combined pathway network (**Fig EV5B**). As for genetic causes of diseases, we further considered a set of 2,661 genes that carried disease-causing mutations (Amberger et al, 2011; Robinson et al, 2008). Specifically, such disease genes were enriched among control proteins in an increasing number of pathways (**Fig 3C**). We corroborated this result by analyzing 11,002 disease genes that were identified from GWAS studies (Hindorff et al, 2009). Furthermore, we obtained similar results when we analyzed the enrichment of such genes in the set of control proteins in the combined pathway network (**Fig EV5C**). Notably, enrichment levels of GWAS related genes were generally lower than genes that harbored disease-causing mutations. Investigating the transformation of a disease to a healthy state, we utilized a set of 2,289 drug targets that were approved by the Food and Drug Administration (FDA) (Knox et al, 2011). Notably, drug targets predominantly appeared in groups of proteins that controlled an increasing number of pathways (**Fig 3D)**. Furthermore, we considered a set of proteins deemed druggable as they carried protein folds, favoring interactions with chemical compounds. Generally, we observed that drug targets were enriched with proteins that controlled many pathways. However, a subset of druggable genes that were not approved drug targets were found diluted. While the enrichment of drug targets in the set of proteins that controlled the combined pathway network was confirmed we found the opposite for both sets of druggable genes that mostly appeared among non-control proteins (**Fig 3E**). To corroborate the independence of these observations, we obtained similar results with Reactome pathway data when we considered the enrichment of disease genes and drug targets among proteins that controlled a large number of pathways (**Fig EV6**) as well as the combined pathway interaction network (**Fig EV7)**.

**Figure 3.**
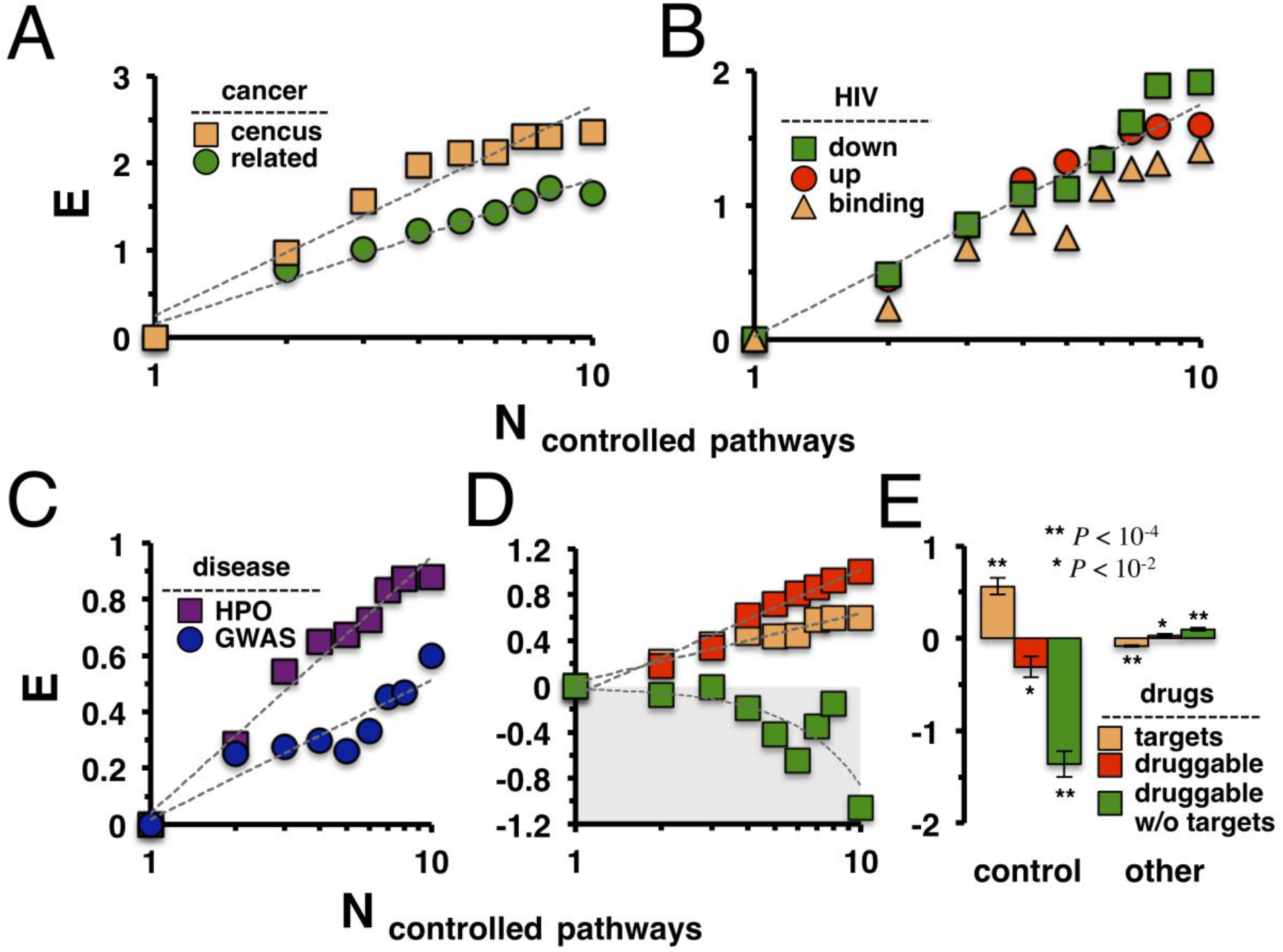
Pathway controlling proteins were enriched with disease genes and drug targets. **A** Using a compilation of census cancer genes and a set of onco- and tumor-suppressor genes, we found that such cancer related genes strongly appeared in groups of proteins that controlled an increasing number of pathways. **B** Such control protein groups were enriched with targets that the HIV virus binds as well as genes that were dys-regulated after viral infection. **C** More generally, disease genes from genetic (HPO) and genomic (GWAS) sources were enriched in groups of proteins that frequently controlled pathways. **D** FDA approved drug targets and druggable genes were enriched in such groups of control proteins as well. However, a subset of druggable genes that excluded known drug targets appeared diluted. **E** In turn, approved drug targets were enriched in the set of proteins that controlled the network of the combined pathway network while druggable genes in general however appeared diluted.

Based on the obtained results so far, we further assumed that the topological and biological role of control proteins may be reflected by their propensity to be evolutionarily conserved. In particular, we labeled all proteins in different organisms that had a human ortholog based on KEGG orthology groups. In the heatmap in **Fig EV8** we however found that proteins that controlled the combined KEGG human pathway network appeared randomly scattered among orthologous proteins in closely related organisms. Constructing directed interaction networks by combining all KEGG pathways of a given organism, we determined control proteins in these combined organism specific networks. In the heatmap in **Fig 4A** we labeled all human proteins with conserved control proteins in closely related organisms. Notably, human control proteins in the combined pathway network significantly aligned with their conserved counterparts in different organisms, an observation that we quantitatively confirmed by a Fisher’s exact test (P < 10^−5^). Extending such considerations, we observed that proteins that controlled an increasing number of human pathways were preferably conserved as proteins that controlled combined pathway networks in different organisms (**Fig 4B**). Notably, enrichment levels differed between *S. scrofa* and *C. familiaris* and more distantly related organisms such as *G. gallus* and *X. laevis*. As a corollary, we hypothesized that human proteins that control an increasing number of pathways have conserved counterparts that control a similar number of pathways in other organisms. Determining control proteins in pathways of different organism separately, we indeed found that the number of pathways that proteins controlled in different organisms correlated well with their human counterparts, indicating different levels of evolutionary kinship (**Fig 4C**).

**Figure 4.**
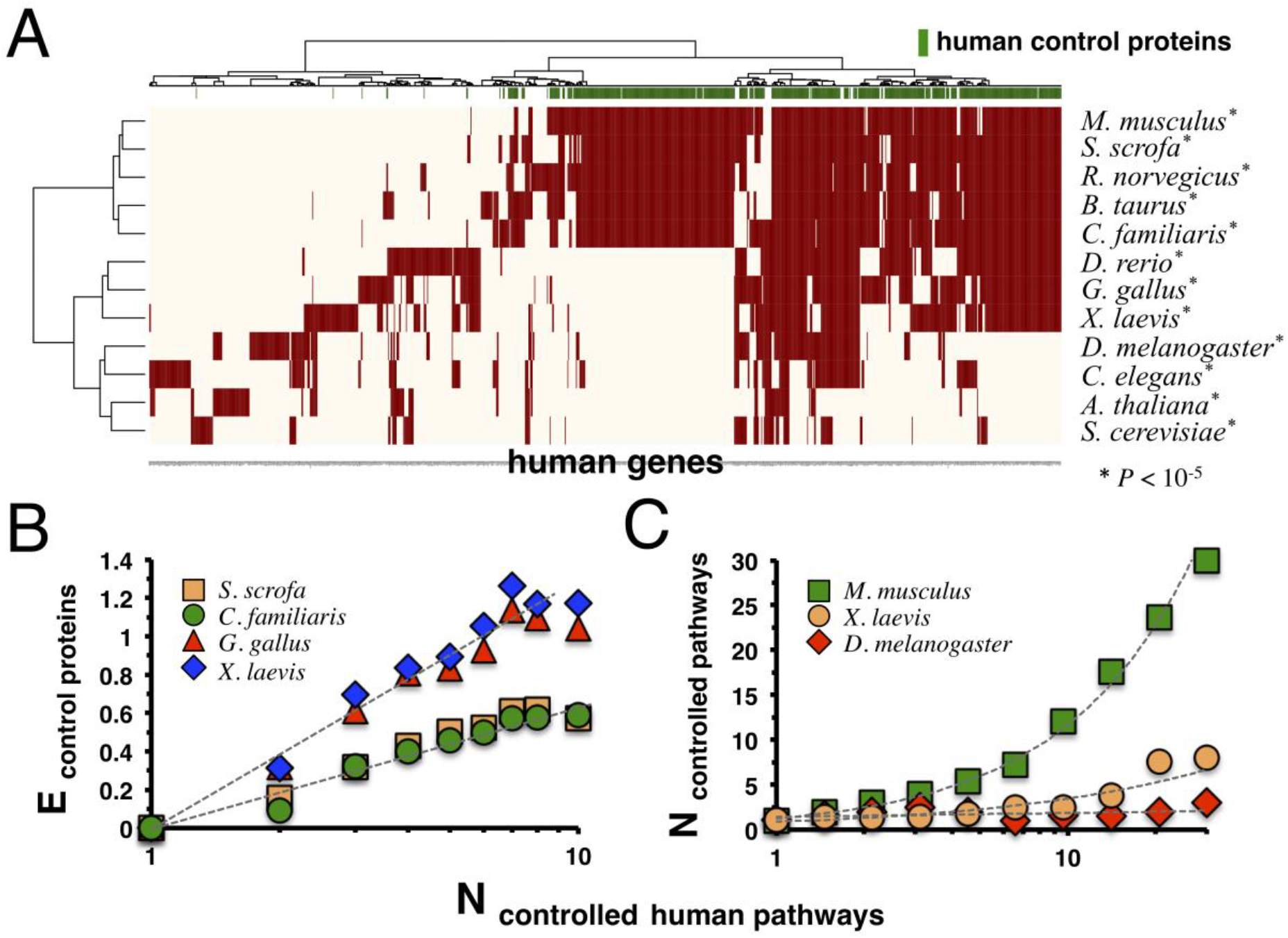
Evolutionary conservation of pathway controlling proteins. **A** We mapped human genes to conserved proteins in different organisms that controlled networks of interactions when we combined all pathway interactions in the underlying organism. Notably, we observed that human control proteins were significantly enriched with corresponding control proteins in other organisms (P < 10^−5^, Fisher’s exact test). **B** Human proteins that controlled an increasing number of pathways appeared enriched with evolutionarily conserved proteins that controlled networks of combined pathways in other organisms. **C** As a corollary, we found that the numbers of pathways human proteins controlled correlated well with their organism specific counterparts.

## DISCUSSION

In this work, we determined proteins that control pathways represented as separate networks of directed molecular interactions. Notably, the frequency distribution of the number of pathways that were controlled by given proteins decayed as a power-law, suggesting a minority of proteins that controlled many pathways and *vice versa*. Such a characteristic may be rooted in the underlying propensity of pathways to substantially overlap, as pathways share genes, allowing pathways to cross-talk. Furthermore, we found that proteins that controlled many different pathways separately also had a heightened chance to control a network of interactions obtained by pooling all pathway interactions. In a similar vein, control nodes that remained robust in networks where we flipped and rewired interactions were preferably enriched among control nodes in an increasing number of pathways. Such observations suggest that the combined pathway network still carried the blueprint of the underlying pathways, transcending different levels of topological organization despite omitting their boundaries.

As such results highlighted the topological placement of control proteins, the question remained if these characteristics translated into a governing, meaningful biological role. Emphasizing their biological importance, control proteins on both topological levels were enriched with essential genes. While signaling proteins were found enriched as well, we observed the opposite for membrane-bound receptor proteins. Such results indicated that pathways were rather controlled by proteins deeply embedded in signaling pathways than by their entry points. In a similar vein, the placement of control proteins may support functional interactions that exert biological control. Indeed, kinases were strongly enriched in sets of proteins that controlled an increasing number of pathways. Furthermore, we observed a lower enrichment of transcription factors, indicating their ubiquitous presence in terms of gene regulation, while kinases may control a large number of pathways to collect and disseminate biological information. As a corollary of this hypothesis, we expected that recipients of post-translational modifications may be control proteins as well. Indeed, we found that acetylated and methylated substrates were strongly enriched in the sets of proteins that controlled an increasing number of pathways. While still enriched, we found much weaker signals when we considered phosphorylated proteins. The latter observation may be a consequence of the fact that a considerable amount of known pathways cover signaling functions that strongly feature phosphorylation events. Nonetheless, the prevalence of kinases and proteins with posttranslational modifications suggested that the placement of control proteins in different pathways was crucial for the dissemination of biological information.

In terms of network medicine, mutations that cause diseases mediate their influence through the interactions an afflicted protein is involved in (Barabasi et al, 2011). As a consequence, we expected that proteins that controlled an increasing number of pathways may be enriched with disease genes as the topological placement of control proteins allows fast transmission of a genomic perturbation. Indeed, disease genes that carried mutations were strongly enriched in sets of control proteins. Surprisingly, we observed that disease genes that were identified from genome-wide association studies were less strongly enriched, most likely reflecting the fact that GWAS identify genomic regions but not specific disease causing genes (Vinayagam et al, 2016). As a corollary, we corroborated the role of control proteins as central to the dissemination of biological information to transform a cell from a disease to a healthy state, when we investigated the enrichment of drug targets. While we found that drug targets and druggable genes were enriched with proteins that controlled an increasing number of pathways, we observed the opposite when we considered druggable genes that were not confirmed as FDA approved drug targets. Considering control proteins in the combined pathway network, we surprisingly found that only approved drug targets were enriched, suggesting that protein domain-folds that can interact with drugs alone are putatively no good indicators of a potential drug target.

As a final consideration, we expected that the topological and biological relevance of proteins that controlled pathways was an evolutionarily conserved feature. Initially, we surprisingly found that control proteins in the human combined pathway network did not appear as particularly conserved in other organisms. Yet, we observed a strong conservation signal, when we considered ortholog proteins that controlled the combined pathway network in different organisms. Transcending different topological levels, human proteins that controlled an increasing number of pathways had conserved counterparts in organism-specific combined pathway networks that well reflected evolutionary distance by corresponding enrichment levels. In particular, increasing evolutionary distance to human was reflected by a decreasing pool of orthologs that may translate into lower enrichment of human control proteins. Furthermore, we observed strong correlations between the number of controlled pathways, when we compared human control proteins to their conserved counterparts in other organisms. Although pathways in other organisms are mostly inferred through the aid of orthologous proteins, our results still suggested that the evolution of pathways retained topological control features as well.

## METHODS

### Pathway information

We collected interaction information of pathways in different organisms from the KEGG (Ogata et al, 1999) and Reactome (Jupe et al, 2012) databases that as parsed with the graphite R tool (Sales et al, 2012). The vast majority of annotated interactions in these pathways were directed, indicating flow of biological information from *e.g.* a kinase to a substrate. Furthermore, physical interactions between proteins (such as protein-protein interactions) were annotated as undirected. We only accounted for directed interactions and considered pathways with at least 5 directed interactions, resulting in 276 KEGG and 1,192 Reactome pathway specific networks. Based on these data sources, all interactions of pathways were pooled to obtain a directed network of 67,038 interactions between 5,398 proteins using KEGG pathways and 8,084 proteins in 180,020 edges using Reactome pathways.

### Functional sets of genes

We collected 2,708 human essential genes from the online gene essentiality database (OGEE) (Chen et al, 2012) and the Database of Essential genes (DEG) (Luo et al, 2014). As for human transcription factors and kinases, we utilized a set of 1,471 manually curated sequence-specific DNA-binding transcription factors (Vaquerizas et al, 2009; Wilson et al, 2008) and 501 genes from the Kinome NetworkX database (Cheng et al, 2014) that collects kinase information from the literature and other databases. As for posttranslational modifications (PTM) we used 17,511 phosphorylated proteins, 6,928 acetylated proteins and 5,418 methylated proteins from the PhosphoSitePlus database (Hornbeck et al, 2015). As for signaling genes, we collected 4,408 genes that were annotated with a signaling function without receptor domain function from Gene Ontology (GO) (Ashburner et al, 2000). Furthermore, we used 5,701 genes that carried a trans-membrane protein domain (Almen et al, 2009).

### Disease genes and drug targets

As representative sets of cancer genes, we used 568 genes that were annotated by the Sanger Center as causally implicated in oncogenesis (Futreal et al, 2004) as well as 1,259 onco- and tumorsuppressor genes that were predicted as cancer-related (Higgins et al, 2007). As for viral infections, we utilized 544 human proteins that interacted with proteins of the HIV virus from the Human Immunodeficiency Virus Type 1 (HIV-1) Human Interaction Database (Ako-Adjei et al, 2015). From the same source we used 788 human genes that were down-regulated and 1,118 genes that were up-regulated upon HIV infection.

As for disease genes, we accounted for 2,661 genes that were identified as causal for a disease as of human phenotype ontology database (HPO) (Robinson et al, 2008) that is based on the Online Mendelian Inheritance in Man (OMIM) database (Amberger et al, 2011). Furthermore, we collected 11,002 disease genes that were identified from GWAS studies (Hindorff et al, 2009).

As for drug targets, we used a set of 2,289 drug targets that were approved by the Food and Drug Administration (FDA) as of the DrugBank database (Knox et al, 2011). Furthermore, we accounted for 2,436 genes that were annotated as druggable as these proteins carried domains that were deemed suitable to interact with drugs (Hopkins & Groom, 2002).

### Controllability analysis

Driver nodes were determined in pathways that were represented as directed un-weighted networks of interactions between genes in the underlying pathways. Such drivers are defined as nodes that are sufficient to ensure the structural controllability of linear dynamics (Liu et al, 2011). In particular, such a structural controllability problem can be mapped to a maximum matching problem, assuming that a network of direct interactions is a graph-based proxy of the underlying dynamical system. The maximum matching problem can be solved in polynomial time by the Hopcroft-Karp algorithm (Hopcroft & Karp, 1974), mapping a directed to a bipartite network. Specifically, we mapped directed links to edges between partitions of nodes that start and end edges. In the matching, a subset of edges *M* is a matching of maximum cardinality in a directed network if no two edges in *M* share a common starting and ending vertex. Vertices that do not appear in *M* are unmatched and have been shown to be nodes that structurally control the underlying network (Liu et al, 2011). As a corollary, a maximum matching implies the presence of a minimum set of such driver nodes of size *N*_*D*_.

To assess the impact of network nodes on the controllability of the underlying directed network we applied the following heuristic (Vinayagam et al, 2016) (**Fig. 1A**): After a node is removed from the underlying network, we determined the size 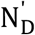 of driver nodes in the changed network. If 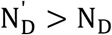, the node is classified as indispensable (*i.e.* a control node) as the number of driver nodes increased. In other words, the deletion of a node increased the number of nodes that allow the control the underlying network. If 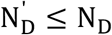, the node is classified as non-controlling as the number of driver nodes remained unchanged or decreased (Vinayagam et al, 2016).

### Enrichment Analysis

In a group *i* of control proteins the corresponding number of proteins with a certain characteristic 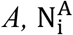 (e.g. being essential or a drug target) were determined. Randomly sampling a set of proteins with characteristic *A*, we calculated the corresponding random number of control proteins with 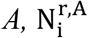. We defined the enrichment of proteins with characteristic *A* in a group *i* of control proteins that appear in a given number of pathways as 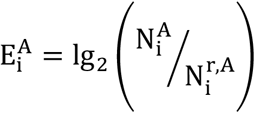

Furthermore, the enrichment of proteins with a certain characteristic *A* was determined as a function of the number of pathways *k*, that given proteins control. In particular, 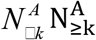 is the number of proteins with *A* that controlled ⁏ *k* pathways. Randomly sampling a set of proteins with characteristic *A*, we calculated the corresponding random number of 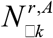. The enrichment of these proteins in a group of proteins that control at least *k* pathways was then defined as 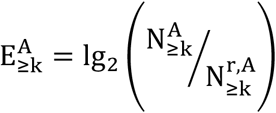. In both cases, 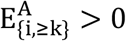 points to an enrichment of feature *A* and *vice versa*. In particular, proteins with feature *A* were sampled 10,000 times and averaged enrichment values thus obtained.

## ACKNOWLEDGEMENTS

Fruitful discussions with M. Afkhami and A. Wilson are greatly appreciated.

